# Signatures of selection in the coral holobiont reveal complex adaptations to inshore environments driven by Holocene climate change

**DOI:** 10.1101/2020.02.25.951905

**Authors:** Ira Cooke, Hua Ying, Sylvain Forêt, Pim Bongaerts, Jan Strugnell, Oleg Simakov, Jia Zhang, Matt A. Field, Mauricio Rodriguez-Lanetty, Sara C. Bell, David G. Bourne, Madeleine JH van Oppen, Mark A. Ragan, David J. Miller

## Abstract

Climate change at the Pleistocene/Holocene boundary reshaped many coastal landscapes, and provides an opportunity to study recent adaptive processes in marine species and ecosystems including coral reefs. On the Great Barrier Reef (GBR) sea level rise flooded a vast shelf creating a distinct inshore region which now harbours extensive coral assemblages despite being subject to relatively high turbidity, freshwater input and thermal fluctuations. To investigate how the coral holobiont has adapted to these conditions we first generated a highly contiguous genome assembly for *Acropora tenuis* based on long-read sequencing, and then used shallow whole-genome resequencing of 148 *Acropora tenuis* colonies from five inshore locations to model demographic history, identify signatures of selection and profile symbiont communities. We show that corals from Magnetic Island, located in the central inshore region of the GBR, are genetically distinct from those 50-500km further north, reflecting a Pleistocene (250-600Kya) split, whereas photosymbiont genotypes differ between reefs in a pattern more likely to reflect contemporary (Holocene) conditions. We also identified loci in the coral host genome with signatures of positive selection in the northern population and used coalescent simulations to show that these are unlikely to be accounted for by demographic history. Genes at these loci have roles in a diverse range of processes that includes heterotrophic nutrition, osmotic regulation, skeletal development and the establishment and maintenance of symbiosis. Our results show that, in the case of *A. tenuis* holobionts from the inshore GBR, the genomes of both the coral host and the primary photosymbiont of have been significantly shaped by their environment and illustrate the complexity of adaptations that have occurred in response to past climate change.

## Introduction

Coastal marine ecosystems worldwide are threatened by the effects of climate change including warming^1,2^, acidification, altered patterns of freshwater input and changing species interactions due to range-shifts^3^. Coral reefs are especially vulnerable because their structural integrity depends on the symbiosis between corals and photosymbionts (family Symbiodiniaceae^4^), but this relationship breaks down under stress, leading to bleaching and widespread mortality during marine heatwaves^5,6^. The projected severity of these impacts therefore depends on the ability of coral holobionts (coral, photosymbionts and associated bacteria^7,8^) to adapt fast enough to maintain the integrity of reef ecosystems^9,10^. A key factor that might improve the adaptive outlook for corals is the existence of stress-tolerant populations that could fuel adaptation in conspecifics through natural or assisted gene flow^11–14^. Ironically, many of these resilient coral populations are threatened because they are found in inshore environments^13,15,16^ where they face additional pressures from reduced water quality^17–19^ that can exacerbate susceptibility to elevated temperature^20–22^.

Although several studies have identified genetic correlates of the thermal stress response or divergent loci that characterise thermal resilience^12,23,24^, little is known about adaptive variation related to other traits that may be important for survival under different local environmental conditions. A comprehensive understanding of local adaptation is important since differences in local fitness and response to stress^13,15,25,26^ can interact with gene flow to produce migrational load^27^ and because locally adapted traits include those likely to be required to survive climate change. One way to understand the genetic architecture and longer-term adaptive responses of corals is to examine signatures of past selection, but this is challenging, because selection can be difficult to distinguish from demographic factors such as population bottlenecks. Here we address this issue using dense allele-frequency information from whole-genome sequencing to detect signatures of selection while using explicit demographic modelling to calculate a neutral distribution and control for false positives. An additional benefit to the whole-genome sequencing approach used here is that it also permits simultaneous genotyping of the dominant photosymbiont in each coral colony.

We apply our approach to identify signatures of selection and profile symbiont communities in *Acropora tenuis* populations from inshore reefs of the GBR. This species is found throughout the Indo-Pacific and is widespread across the GBR where it is common on both inshore and offshore reefs^28,29^. The species has also become an important model for research into water-quality stress^30,31^, adaptive changes in symbiont composition^32^ and the feasibility of assisted evolution approaches^33,34^. The reefs sampled in this study provide an ideal context in which to study local adaptation because they were recolonised between 14 and 6Kya when rising sea levels resulted in marine flooding of the GBR shelf^35^. This event changed the environment available for coral growth dramatically, shifting from a narrow line of fringing reefs at the edge of the shelf to the distinct outer and inner reefs characteristic of the GBR today^35^. Water quality on GBR is a major driver of coral biodiversity and differs markedly between inshore and outer reefs^36^, with inshore sites typically having much lower water clarity and higher nutrient levels^36^. Water quality depends on many factors, some of which, such as the frequency of exposure to flood plumes varies significantly between inshore locations^37,38^. Our results highlight the complex molecular changes associated with corals that have colonised these inshore habitats. In addition, we investigate differences between inshore sites with contrasting water quality due to relative proximity to large river systems^39^ and find that they host distinct populations of photosymbionts.

## Results

### *De-novo* assembly of the *Acropora tenuis* genome

To maximise our ability to model demographic history and detect selective sweeps, we generated a highly contiguous and complete reference genome for *Acropora tenuis* based on PacBio long-read sequencing. The resulting assembly of 487Mb in 614 contigs with an N50 of 2.84Mb, and largest contig of 13.5Mb, represents a considerable improvement in contiguity compared with most previously published coral genomes (Supp Table S1) including a recent assembly from the same species^40^. Importantly, over 80% of the genome assembled into contigs larger than 1Mb, allowing us to minimise artefacts arising from ambiguous alignments from unmerged haplotypes or misassemblies at the ends of contigs. Large genomic fragments also provide the necessary length to capture recent selective sweeps which can distort allele frequencies over centimorgan-sized regions^41,42^.

Annotation of this genome assembly with Illumina RNA-seq data resulted in 30,327 gene models of which over 90% (27,825) belonged to one of 14,812 ortholog gene family groups shared with other cnidarian genomes. Of these gene families, 97% (14,403) were shared with other published *Acropora* genomes (*A. millepora* and *A. digitifera*). BUSCO analysis identified a near complete 91.8% set of core metazoan genes including a small fraction that were duplicated (2.6%) or fragmented (1.8%). A genome size estimate of 470Mb obtained from kmer analysis of short read data with sga.preqc^43^ was close to the final assembly size, confirming the high completeness of the assembly and suggesting that there are few artificial duplications remaining after purging of unmerged haplotypes.

The completeness of our long-read assembly allowed us to resolve a substantially larger proportion of interspersed repeats (41.7%) compared with the published acroporid assemblies based on short-read data (*A. digitifera^44^*: (30.2%), *A. millepora^45^* (31.6%), *A. tenuis^40^* (34.3%). The only other available coral genome with comparable repeat resolution is a recently released long-read assembly for *A. millepora^46^*, which identified a very similar proportion (40.7%) of interspersed repeats to our *A. tenuis* assembly. This increased ability to resolve repeats is reflected in a substantially higher proportion of unclassified elements in long-read assemblies (20.8% *A. tenuis*; 21.2% *A. millepora*) compared with short-read assemblies (16.4% *A. digitifera*; 16.5% *A. millepora*), but the overall composition of known repeat classes was remarkably similar between *Acropora* genomes (Supp Figure S1).

As an additional check on the accuracy of our *A. tenuis* assembly we compared its high-level genomic organisation with that of the congeneric species, *A. millepora* for which a chromosome-scale linkage map is available^46^. Macrosynteny analysis between the two *Acropora* genomes revealed a high degree of collinearity and limited rearrangements between linkage groups (Figure 1A) which most likely reflects their small evolutionary distance (approximately 15Mya^40^). For comparison we also examined synteny between *A. millepora* and the much more distantly related anthozoan, *Nematostella vectensis* (greater than 500Mya^47^). Despite this long divergence time the number and basic structure of linkage groups was remarkably well maintained, but with near complete rearrangement within groups (Figure 1B).

**Figure 1:**
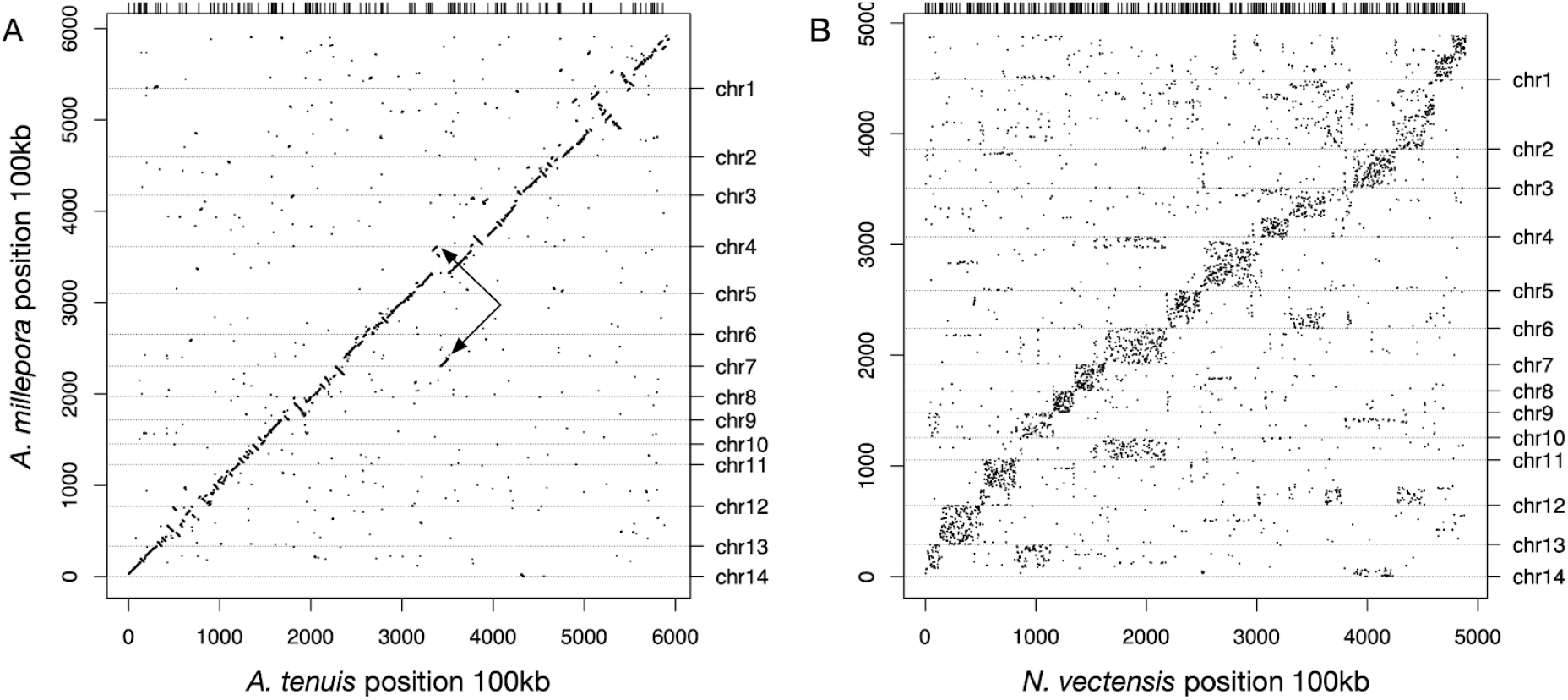
Macrosynteny conservation between *A. tenuis* and *A. millepora* (panel A) and between *Nematostella vectensis* and *A. millepora* (panel B). Dots represent blocks of collinear groups of genes and margin lines represent boundaries between contigs (top) or *Acropora millepora* linkage groups (right). In panel A very few *A. tenuis* scaffolds map across *A. millepora* chromosomes (a single example is shown with arrows). In panel B the blocks along the diagonal reflect ancient conserved linkage groups which roughly correspond to chromosomes in *A. millepora*.

### Population structure

To investigate potential adaptive changes in the *A. tenuis* holobiont in relation to water quality, we chose reefs from opposite ends of two plume influence gradients, Burdekin in the south and Wet Tropics in the north (Figure 2A). These gradients capture long-term frequency of exposure to riverine flood plume waters based on aerial and satellite monitoring data^37,38^ as well as predictive modelling (see methods for full details and rationale). Although other factors are likely to also be involved, flood plume exposure is a strong driver of water quality^48^. For the Burdekin gradient, two sites with high river influence (“plume”; Magnetic Island and Pandora Reef) and one site with low riverine influence (“marine”; Pelorus Island) were sampled, whereas for the Wet Tropics gradient we sampled one “plume” (Dunk Island) and one “marine” site (Fitzroy Island).

**Figure 2:**
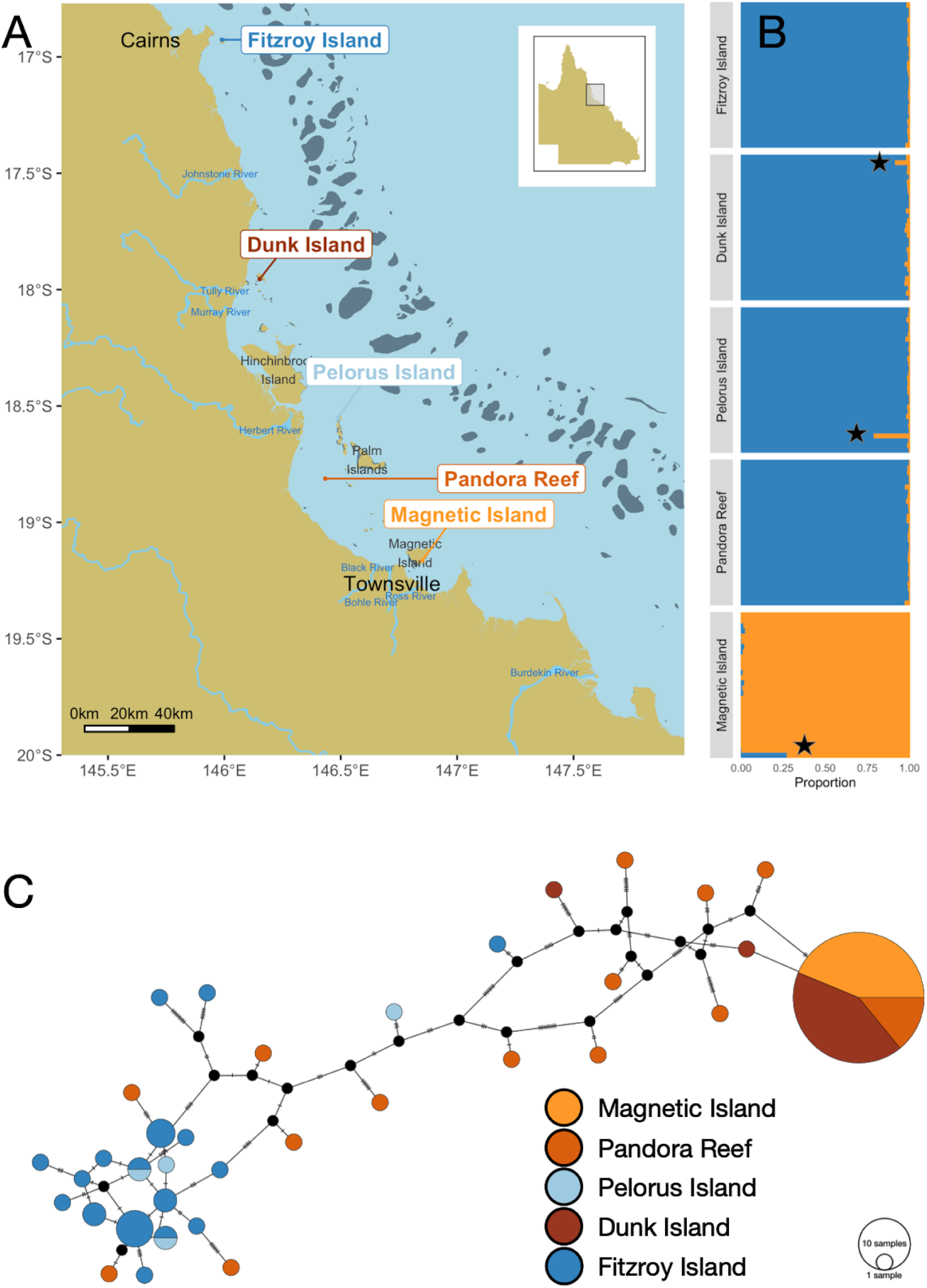
Population structure of coral host and dominant (*Cladocopium*) photosymbionts. Plots A and C use the same colour scheme (bottom right) which shows plume sites in shades of orange and marine sites in shades of blue. **A.** Sampling locations in the central inshore Great Barrier Reef, Australia. Major river systems are labelled and affect sites northward due to the direction of residual inshore current. **B.** Coral host admixture proportions based on NGSAdmix analysis for each of the five sampling locations. The northern population is shown in dark blue while Magnetic island is in orange. Samples indicated with a black star represent likely hybrids. **C.** Mitochondrial haplotype network based on the dominant strain of the symbiont *Cladocopium* in each sample. Substitutions separating nodes are represented by cross bars.

Superimposed on these water-quality gradients, we observed strong coral population structure based on latitude. *Acropora tenuis* colonies from Magnetic Island were genetically distinct from those inhabiting the four reefs north of Magnetic Island. Analysis of population structure with PCAngsd showed these as two clear clusters (Supp Figure S2) and admixture analysis showed near complete assignment to either of these clusters for all but three individuals (Figure 2B). We refer to these clusters as north (all reefs other than Magnetic Island) and Magnetic Island. Genome-wide average Fst values between Magnetic Island and northern reefs were much larger (0.19) than between reefs within the northern cluster (0.008). We also observed evidence of recent gene flow between Magnetic Island and northern reefs with three colonies showing mixed ancestry (Figure 2B). We also checked to determine if this population structure was captured by differences in mitochondrial haplotypes between colonies but found that, in contrast to the nuclear data, mitochondrial haplotypes were completely unstructured (Supp Figure S3). This most likely reflects the relatively recent divergence (see below) combined with a slow mutation rate which is a well-documented feature of anthozoan mitochondrial genomes^49–51^.

### Demographic history

To model demographic history and interpret and control for error in selection analyses, the complementary approaches MSMC and ∂a∂i were used, with combined results from both suggesting that the Magnetic Island and northern populations diverged between 600Kya and 270Kya (Figure 3A and 3B). Estimates using ∂a∂i are based on the best-fitting model (“isolation_asym_mig”), which allows for a period of isolation followed by secondary contact with asymmetric migration (See Supp Table 2 for full parameter lists of all models). An independent estimate based on divergence of population size trajectories inferred by MSMC suggests a slightly older split, with first divergence occurring at around 600Kya and becoming clearly distinct by around 300Kya. Both ∂a∂i and MSMC models also suggest that inshore populations of *A. tenuis* have recently declined or have become disconnected from larger parent populations. This is particularly pronounced in the case of the Magnetic Island population, which both modelling approaches estimate as having larger historical and much smaller contemporary effective population sizes (Figure 3: also see all alternative ∂a∂i models in Supp Figure S5). Although this bottleneck was consistently observed across all models, MSMC estimates of the timing were much older (~100Kya) than those inferred by the best-fitting ∂a∂i model (3Kya). In this instance ∂a∂i estimates are more likely to be reliable since the single diploid genomes used for MSMC (one each from Magnetic Island and Northern populations) provide few recent coalescent events with which to estimate recent demographic change^52,53^.

**Figure 3:**
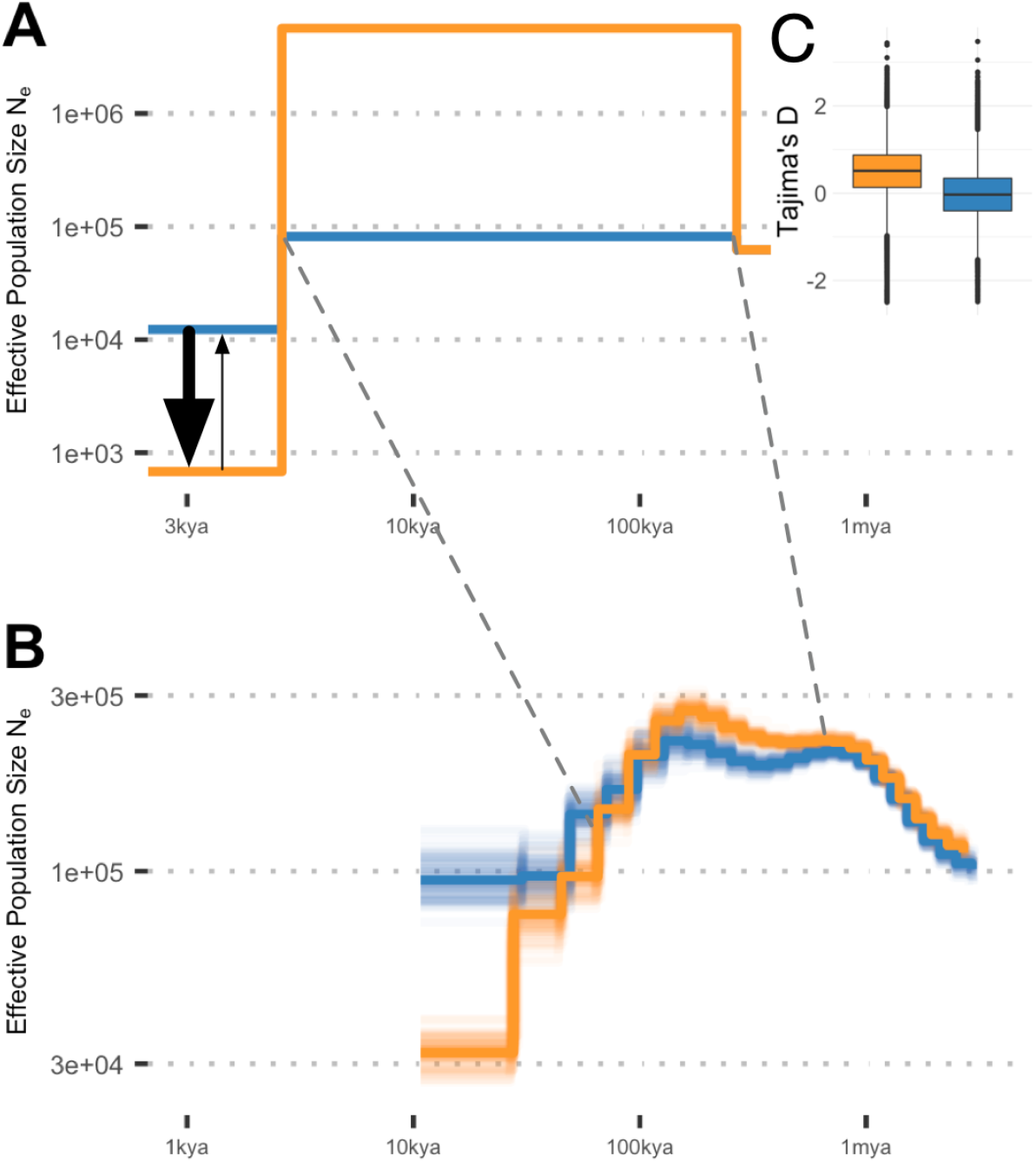
Population genomic statistics and models of demographic history. Orange represents Magnetic Island and blue represents northern populations in all component figures. Dashed lines connect shared events in both models, the point of divergence between populations, and onset of recent bottleneck. **A** Changes in effective population size and estimated gene flow for the best-fitting ∂a∂i model. Relative gene flow is shown using arrows with larger flow from north to south indicated by arrow size. **B** Changes in effective population size estimated using MSMC from deep sequencing of a single colony from Magnetic Island (MI) and a colony from Fitzroy Island (FI). **C** (Inset top right) Genome wide estimates of Tajima’s D calculated using PCAngst.

Genome-wide estimates of Tajima’s D are sensitive to demographic history and were consistent with the main demographic inferences from ∂a∂i and MSMC analyses (Figure 3C). In particular, a high value for Magnetic Island is consistent with a recent or ongoing bottleneck for that population^54^, whereas values close to zero for northern reefs are consistent with minimal bottleneck effects there. Demographic models that included asymmetric migration provided a better fit to the 2D SFS than those without migration or with symmetric migration (Figure3: Supp Table 2). Across all of these models migration coefficients were larger for the southward direction. Although this is against the predominant direction of inshore current flow^55^ and predictions based on biophysical modelling^56^, it is consistent with studies that have shown southward migration to be the predominant direction of gene flow for *A. tenuis* and *A. millepora* on the GBR^9,56^ including for inshore populations. This disconnect between biophysical modelling (which reflects northward inshore currents) and apparent gene flow could reflect differences in effective population size. Demographic modelling and genome-wide Tajima’s D suggest a strong bottleneck and small effective population size for Magnetic Island and, since migration coefficients in ∂a∂i are measured as a proportion of individuals at the destination, the small size of the Magnetic Island population could skew outcomes toward southward flow.

### Photosymbiont differences between reefs

The availability of complete genomes from five major genera within the Symbiodiniaceae allowed us to identify (using kraken^57^) between 100k and 300k reads of symbiont (Symbiodiniaceae^4^) origin in each sample (Supp Figure S4). For almost all samples the dominant symbiont genus was *Cladocopium* (formerly Clade C^4^) with a single sample showing a significant proportion of *Durusdinium* (formerly Clade D^4^). Although the numbers of symbiont reads were insufficient to permit genotyping on nuclear loci, these did provide sufficient depth (at least 5x) for determination of the dominant mitochondrial haplotype in each sample. The resulting haplotype network appeared to be structured according to location, with the majority of samples from “plume” (Magnetic Island, Dunk Island and Pandora Reef) sharing a single haplotype. Samples from “marine” locations (Fitzroy Island, Pelorus Island) did not include this haplotype but instead were highly diverse, forming a sparse network of divergent types. A test for genetic structuring using AMOVA confirmed that differentiation by reef was significant (p<1e^−5^) but it was not possible to link this unambiguously with the water quality designations from our study design (plume vs marine) (p=0.1). Notably, the structuring of Symbiodiniaceae populations by reef followed a biogeographical pattern that was clearly distinct to that of hosts. Although the results point toward long-term frequency of plume exposure as one potential driver of these differences, we cannot rule out other factors such as thermal history or acute differences in plume exposure. Nevertheless, the observation of clear structuring by reef suggests adaptation to local conditions.

### Signatures of selection

A major challenge when identifying genomic sites under selection is that demographic history and migration can produce confounding signals ^58,59^. To estimate false positive rates from such processes, SweepFinder 2 was used in combination with coalescent simulations under a range of demographic models (all ∂a∂i and MSMC models), to generate neutral data from which composite likelihood ratio (CLR) statistics could be calculated. Using this data as a control, we were able to calculate an empirical false discovery rate for a given CLR threshold and neutral model (see methods). For northern reefs we found that, irrespective of the neutral model used, it was always possible to find a CLR threshold that would control false positives to less than 10% (Supp Figure S6). This was not the case for Magnetic Island however, where its extreme demographic history (strong bottleneck) combined with small sample size meant that distinguishing neutral from selective processes was highly model-dependent and uniformly required a higher CLR threshold. We therefore focused our attention on the northern reefs where a CLR threshold of 100 was sufficient to control the false discovery rate to less than 10% under the best fitting model and almost all other demographic models. In support of their status as putative sites under positive selection, the 98 loci passing this threshold also tended to have extremely low values of Tajima’s D (Figure 4) and were often associated with regions of high differentiation (high pairwise Fst) between Magnetic Island and northern reefs (Figure 4). Although SweepFinder 2 is designed to detect classical hard sweeps caused by positive selection, a similar signature can also be produced through persistent background selection^58,60^. Since background selection involves removal of deleterious alleles it should act most strongly on genes with highly conserved functions and is therefore more likely to act similarly in both Magnetic Island and northern populations. Based on this reasoning, 20 loci with low Fst values (<0.1) were removed in order to maintain our focus on genes under positive selection. Finally, a further 7 loci with positive values of Tajima’s D were removed because these could represent genes under balancing selection^54^.

**Figure 4:**
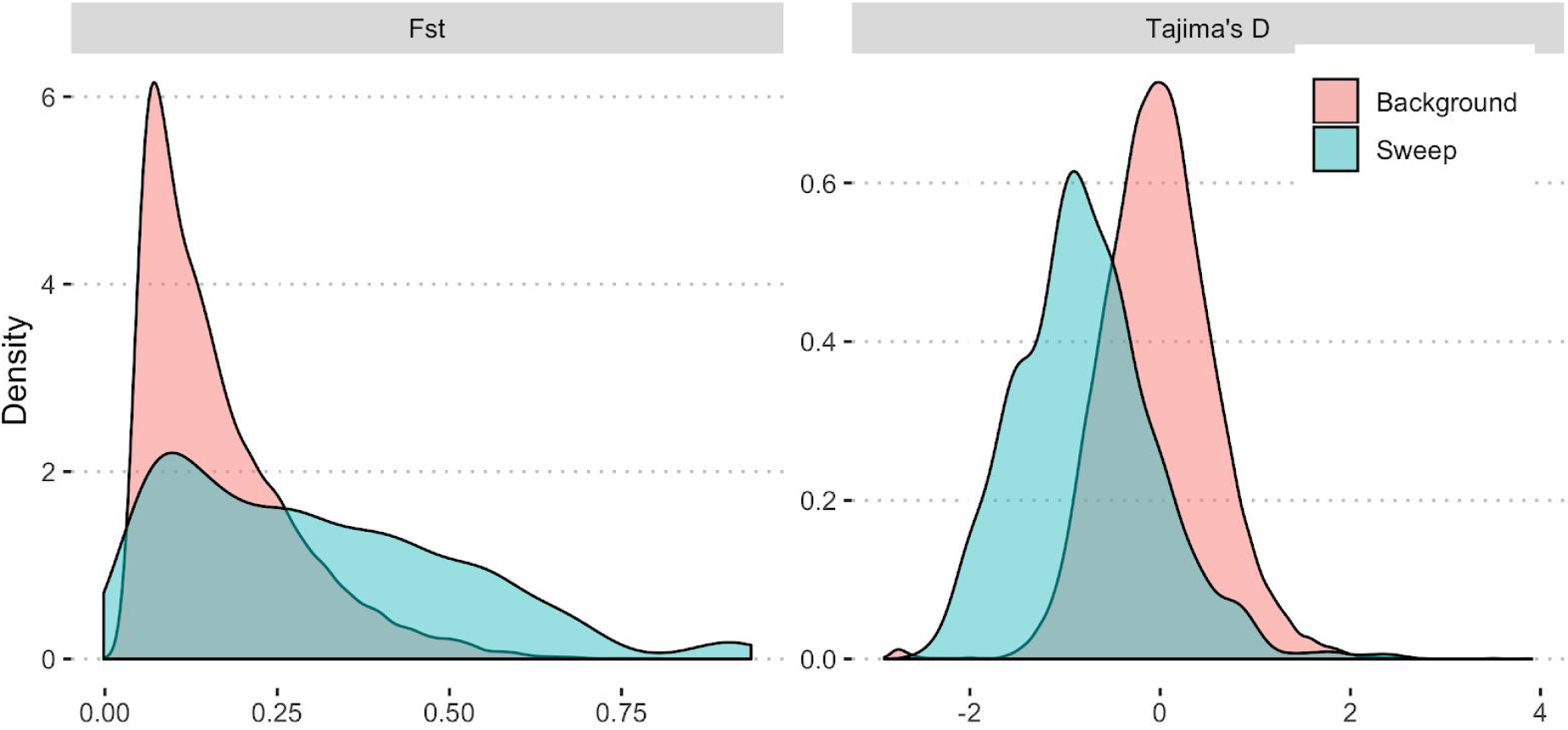
Distribution of genome-wide Fst and Tajima’s D for all loci (red) and regions with CLR > 100 (blue) in the northern reefs population. Horizontal axis shows the relevant statistical value (Fst or Tajima’s D) and vertical axis shows the relative abundance (as normalized density) of loci with the corresponding value.

Our final list of 71 regions impacted by putative selective sweeps were spread throughout the genome (Figure 5) and overlapped with 61 genes, of which 39 could be annotated based on homology or the presence of conserved domains. A complete list of sweep loci, along with associated genes and their functional groupings, is provided in supplementary information.

**Figure 5:**
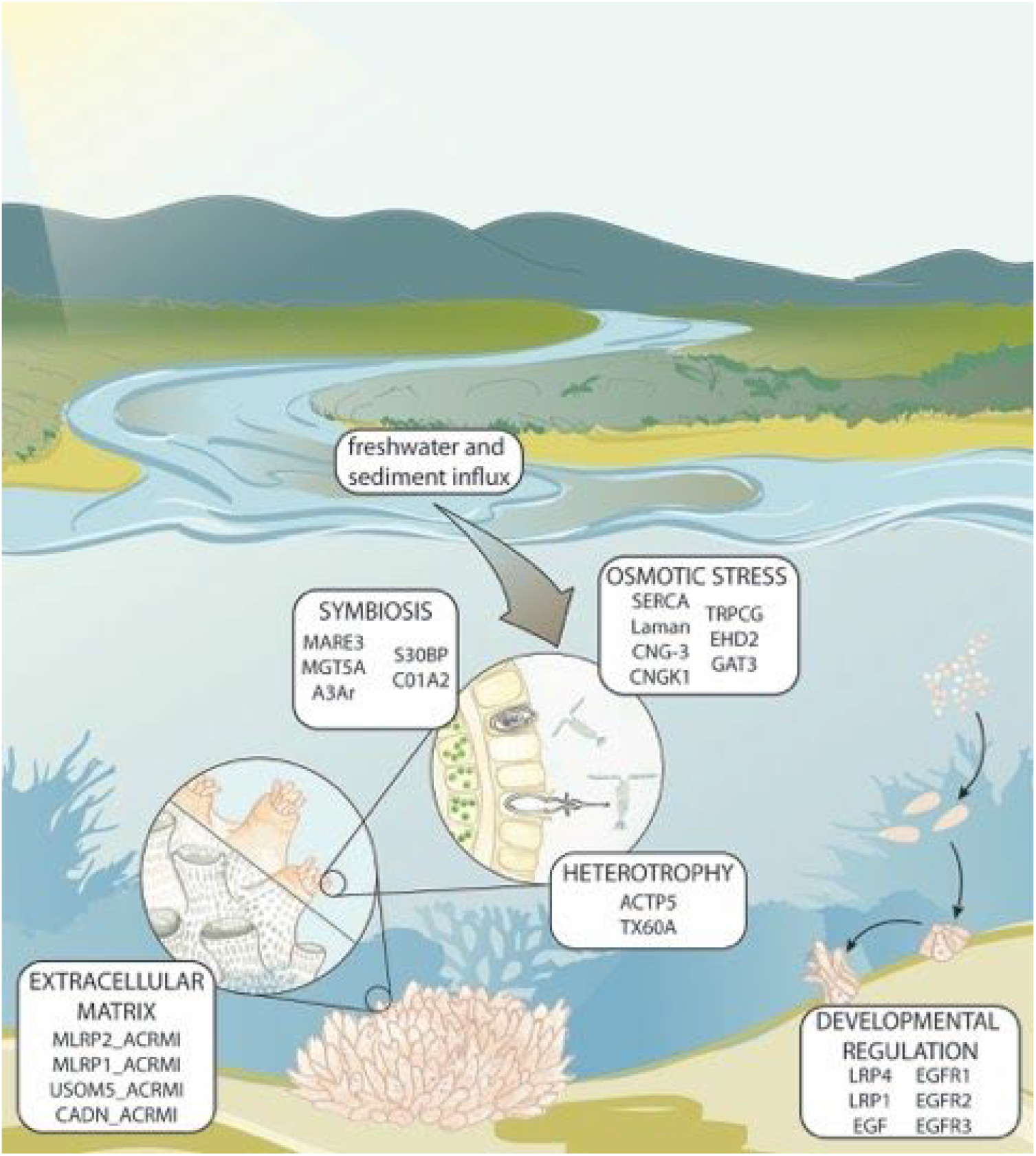
Genes associated with sites under positive selection arranged according to their putative role in adaptations to an inshore environment.

One of the characteristics of selective sweeps is that the signature of selection decays rapidly after fixation has occurred^61^. Even under ideal conditions (no demographic change, classical hard sweep) the CLR has very little power to detect sweeps that originated more than 0.5Ne generations ago and is best able to detect more recent sweeps (0.05-0.2Ne generations ago)^58^. Our best estimate of contemporary Ne for the northern population comes from ∂a∂i simulations (Figure 3) and is around 10000. If this is correct, it puts an upper bound on sweep ages at 25 Kya with peak detectability between 2500 and 10Kya. This timing suggests that many of the sweep loci identified here could represent signatures of selection that arose as a consequence of dramatic coastal change, including creation of the GBR shelf and inshore environment at the start of the Holocene. Even more recent changes, such as increases in sediment and nutrient flux since European settlement ^62,63^, could also produce signatures of selection but our data lack the resolution required to make inferences about this. Key adaptive challenges posed by occupation of the GBR shelf at the start of the Holocene include higher turbidity ^64^, regular influx of riverine sediment and nutrients^19,65^ as well as occasional periods of reduced salinity during severe flooding events ^17,37,38^. Our results suggest that significant adaptation to these conditions has been complex, involving selection on a wide range of genes as well as intergenic loci. Understanding the functional importance of these adaptations is challenging since a significant proportion of loci under selection involve genes that lack homologues in well-studied taxa. This is reflected in the results of Gene Ontology enrichment testing where we searched for GO terms over-represented in sweep genes using topGO^66^. A single, relatively generic term (GO:0005509 calcium ion binding; p = 4.5e^−5^) associated with EGF domain-containing genes and Skeletal Organic Matrix Proteins (SOMPs) was found to be enriched. To complement this formal analysis, we also searched for sweep genes among recent gene-expression and comparative genomic studies in corals and other symbiotic cnidarians. This analysis (detailed below) revealed that many genes associated with sweeps were implicated in roles related to environmental conditions associated with inshore environments of the GBR (Figure 5).

An acute physiological challenge that is particularly important in inshore environments^18^ (compared with offshore) is the increased prevalence of stressful ^67,68^ and potentially lethal ^69,70^ hypo-saline conditions. A recent study using RNASeq found that genes involved in solute transport, amino-acid metabolism and protein recycling were major components of the response to osmotic stress in *Acropora millepora ^71^*. Our results suggest that inshore corals have experienced selection associated with genomic regions close to many of these genes. This includes homologs of the calcium and potassium ion transporters SERCA, CNG-3, CNGK1, TRPCG and GAT-3, EHD2 which regulates vesicle transport, and Lysosomal alpha-mannosidase (Laman) which is involved in glycoprotein recycling. Notably three of these (SERCA, GAT-3, Laman) were very close homologs (97, 82, 97 percent identity respectively) to proteins differentially expressed in response to osmotic stress in *A. millepora*^71^.

Structuring of symbiont genotypes according to local conditions on individual reefs and possibly to water quality (Figure 2C) suggests that the ability to efficiently interact with specific symbionts may be a key requirement for adaptation of the coral holobiont. In support of this, we observed 5 genes associated with selective sweeps in the host genome that were implicated in the establishment and/or maintenance of symbiosis. All of these were homologous to genes observed to be differentially expressed in symbiotic vs aposymbiotic states in *Acropora digitifera* larvae^72^, adult corallimorpharians^73^ or *Aiptasia^74^*, an anemone model for symbiosis. This group included genes involved in the cytoskeletal remodelling events required for symbiosome formation (MARE3, CO1A2), as well as apoptotic regulators (A3Ar, S30BP). In addition, a larger group of genes under positive selection included those with dual roles in symbiosis and developmental regulation such as low-density lipoprotein receptors^74^ and a variety of EGF domain-containing proteins.

While EGF domain-containing proteins are common in animal genomes, a specific class of these (here called EGFRs) containing large tandem arrays of calcium binding type EGF domains appeared to be under strong selection in *A. tenuis*. Two genes encoding EGFRs were associated with exceptionally strong sweep signals located adjacent to each other on a single scaffold of the *A. tenuis* genome. In addition a further five EGFRs were located within 200kb of these strong sweep signals (Figure 5). Although the large tandem arrays of EGF domains in these proteins resemble those in the TGF-beta binding proteins Fibrillin and LTBP, they lack the TB domains that characterise the latter class ^75^. Proteins with various numbers of EGF-like domains have been reported as part of the skeletal organic matrix of both *A. millepora* ^76^ and *A. digitifera* ^77^, but developmental gene-expression data suggest that they may have a more general role in cell adhesion, which then leads to their incorporation in the skeletal organic matrix ^77^. Another potential role for these proteins is in the establishment and maintenance of symbiosis, with separate studies on *Hydra* ^78^ and corallimorpharians ^73^ both reporting increased expression in the symbiont infected state compared to aposymbiotic controls.

**Figure 5:**
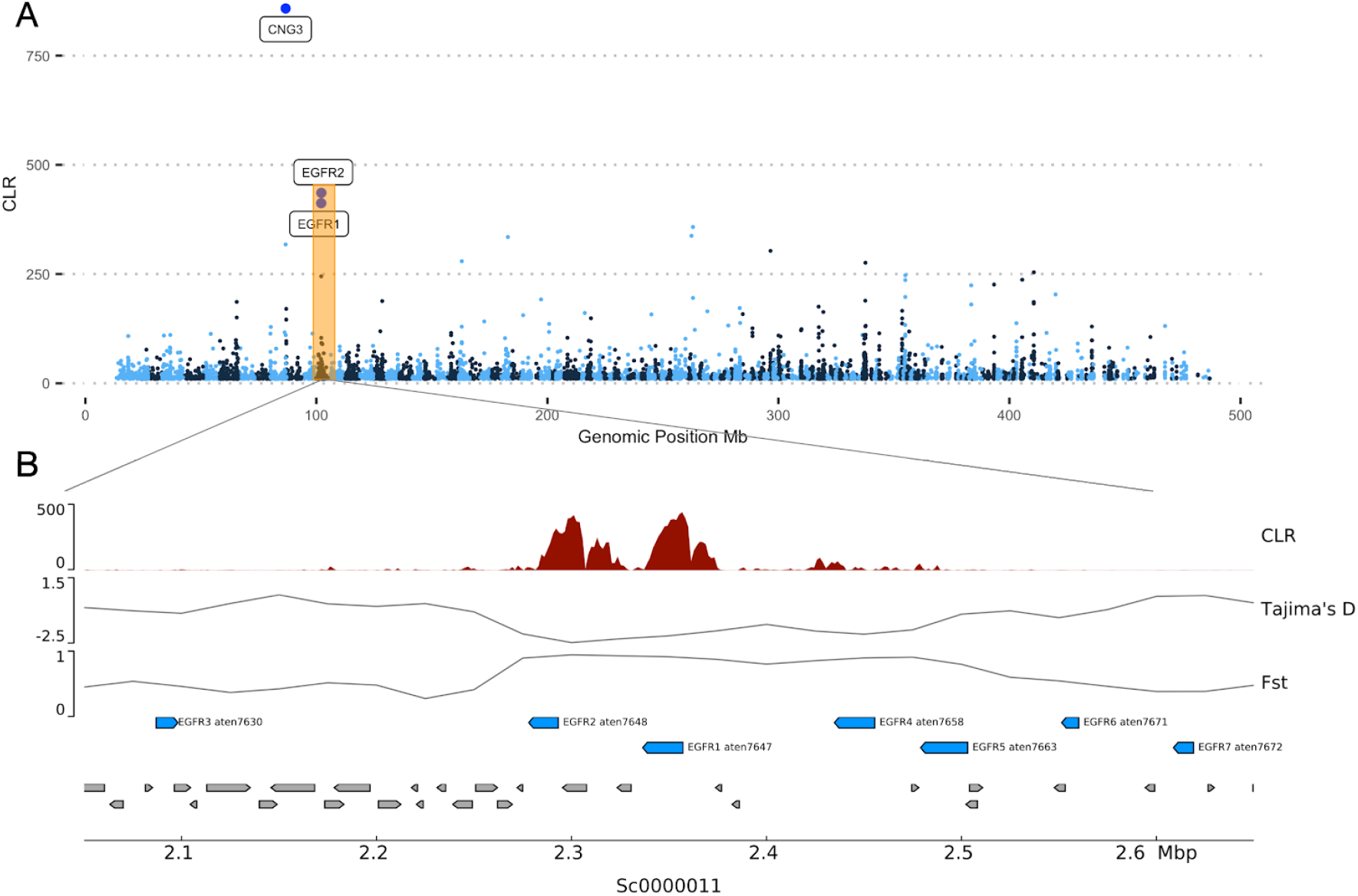
Sites under selection. A Manhattan plot showing genome wide distribution of the SweepFinder 2 CLR statistic. B Detailed view showing a region of the *A. tenuis* genome with evidence of strong selection on two genes encoding EGF repeats (EGFR1, EGFR2). These, and other EGF repeat-containing genes surrounding the sweep are numbered and shown in blue. All genes other than EGF repeat-containing genes are shown in grey.

For several coral species, higher rates of heterotrophy have been observed on inshore reefs relative to offshore reefs^79,80^ and similar effects have also been experimentally demonstrated to occur in response to increased turbidity^81^. In addition, experiments on *Acropora millepora* and *Pocillopora damicornis* showed increased heterotrophic feeding efficiency in colonies from inshore reefs compared with conspecifics sourced from offshore ^80^. Since cnidarians - including corals - rely on toxins injected into prey to assist with heterotrophic feeding ^82^, an important aspect of this inshore adaptation may involve modifications to the toxin repertoire. In support of this hypothesis, we observed strong selection on two homologs of anemone pore-forming toxins, Delta actitoxin (aten_0.1.m1.22362) and Delta thalatoxin (aten_0.1.m1.29056). Delta thalatoxin is the major toxin component isolated from nematocysts of the Okinawan sea anemone (*Actineria villosa*)^83^ and both toxins have strong haemolytic activity ^83,84^ which is a feature of coral-specific toxins identified so far ^85^.

Proteins involved in developmental regulation and formation of the skeletal organic matrix constituted a large fraction of genes under selection. Genes in this category included four of the 36 SOMPs originally identified through proteomic analysis of *Acropora millepora* skeletal materials^76^. This observed enrichment for proteins that potentially affect growth and morphology is consistent with reports of major differences in skeletal growth rates and densities for *A. tenuis* along a water-quality gradient^39^. A natural extension of this, which has been demonstrated in other species,^86^ is that these differences should also exist between inshore and offshore environments. Our results suggest that skeletal organic matrix proteins may play an important role in the mechanism of this morphological adaptation.

### Differential selection between marine and plume environments

Although all four northern reefs essentially form a single panmictic population, strong local selection under low gene flow can lead to differentiation at specific loci^24^ which could manifest as independent soft-sweeps in different locations. To search for evidence of local adaptation due to differences in water quality we performed SweepFinder 2 analyses separately on each of the four northern reefs and identified sweeps that were present exclusively in plume or marine habitats. To accommodate smaller sample sizes when analysing each reef independently as well as reduced signal strength due to ongoing gene flow between these reefs we used a slightly relaxed CLR threshold of 50 (FDR < 15%). This revealed seven sweeps that were exclusive to plume sites and eight exclusive to marine (Supp Info). Several genes in both locations had functions related to transcription, probably reflecting differential expression of as-yet unknown genes. In addition, genes associated with sweeps in plume included a Mucin-like protein (MLP_ACRMI) which forms part of the skeletal organic matrix in *Acropora^76^*. Local adaptations related to skeletal organic matrix proteins in plume locations are particularly interesting in light of transplant experiments which showed dramatic skeleton loss when corals were transplanted from a plume (Magnetic Island) to a marine location (Pelorus Island)^26^. Another interesting protein under selection in plume locations was a homolog of NLRC3, which is an important modulator of Toll-like receptors in humans and part of the NLR (nod-like receptor) family. In corals NLR family proteins have been shown to be downregulated during establishment of symbiosis^87^ and strong selection on NLRs might therefore be involved in maintaining the high symbiont specificity seen in these reefs. Sweeps exclusive to marine locations included a particularly strong signal associated with a gene encoding a glutaredoxin domain containing protein, GRCR2. Proteins in this family are upregulated during heat stress in *Acropora palmata^88^* and have been found to be under positive selection in *Acropora^89^*.

## Discussion

The survival of inshore coral reefs under climate change depends on their ability not only to adapt to increased thermal stress but also to deal with potential changes in precipitation and drastic changes in water quality that can accompany sea level rise^35,64^ as a result of coastal erosion and resuspension of terrigenous sediments. Our results show that *A. tenuis* holobionts from the inshore GBR have been significantly shaped by the inshore environment and fine-scale local differences between reefs on the GBR shelf. The most striking differences at a local scale were found in the photosymbiont association which seemed to be highly specific (single dominant mitochondrial haplotype) at some reefs and much less so (multiple divergent types) at others. While our study points toward water quality as a driver, additional sampling locations coupled with more detailed long-term environmental data are required to understand the ecological basis for this genetic pattern.

The availability of a high-quality reference genome for *A. tenuis* and the use of whole-genome resequencing was crucial to our ability to confidently detect genomic regions under selection and to infer demographic history for the host. Although this approach has rarely been used so far in coral population genetics (although see^90^), the moderate size of many coral genomes (~500Mb) and continually falling sequencing costs means that whole-genome sequencing is now cost competitive with reduced representation approaches for population genomics. Our study demonstrates that the dense allele frequency information afforded by whole genome approaches enables powerful inferences even with relatively shallow sequencing coverage per-genome. As high quality genome assemblies become available for a broader range of corals and their symbionts we expect that this and related approaches will become a key tool in understanding the interaction between past climate change and the evolution of corals and coral reefs.

## Methods

### Reference genome sequencing, assembly and annotation

All genome sequencing was based on DNA extracted from sperm collected from a single *A. tenuis* colony collected on 10th of December 2014 from Orpheus Island.

The species examined in this paper is widely referred to as *Acropora tenuis* on the Great Barrier Reef; however, we note that the taxonomy of this species is likely to change in light of recent molecular phylogenetic research indicating that traditional morphological taxonomy does not accurately reflect species boundaries or evolutionary relationships ^91^. The identity of the species is currently under examination using an integrated taxonomic approach, with preliminary analysis suggesting the species is most likely not *A. tenuis* (Dana 1846) but *A. kenti* (Brook 1892), a species described from the Torres Strait but synonymised with *A. tenuis* (Dana 1848) by Veron and Wallace (1984). Field images and a voucher specimen of the colony used in this study have been deposited in the Queensland Museum (QM Registration Number G335181). After taxonomic revisions are complete, this specimen will be compared to the holotypes of both *A. kenti* (Brook 1892) and *A. tenuis* (Dana 1848), as well as topotypes of specimens from both Fiji and the Great Barrier Reef to determine its actual identity.

An initial round of sequencing was performed on an Illumina HiSeq 2500 instrument using sheared and size selected DNA (~ 450bp) to generate 88.6 million 250bp paired end reads. Further sequencing was then performed on a PacBio RSI instrument by Ramaciotti Center for genomics using 62 SMART Cells resulting in a total of 50.6 Gb of long read data (~108x genome coverage) with read N50 of 15Kb. In addition, to facilitate gene annotation, RNA was extracted from tissues collected from the same colony and sequenced on an illumina HiSeq 2500 instrument. All genome sequencing data are available under ENA accession PRJEB23295.

A draft reference genome was assembled purely from PacBio reads using the FALCON (version 0.7.3) assembler ^92^ followed by separation of haplotigs from primary assembly with FALCON-unzip and error correction with Quiver ^93^. The resulting primary assembly was highly contiguous (N50 ~1320 Kb) but had a larger total size (629Mb) than the haploid size of 470Mb estimated by kmer analysis of Illumina data with sga.preqc ^94^. Mapping of transcript sequences to the draft genome assembly revealed the presence of numerous alternate haplotypes in the primary assembly which most likely accounts for the inflated assembly size. To address this issue we used Haplomerger 2 to identify and merge alternate haplotypes. Finally, small redundant scaffolds less than 2kb in length were identified by BLAST and removed. This resulted in a final merged assembly with a primary size of 487Mb and N50 of 2837 Kb. As any unmerged haplotypes that might remain should be located entirely on the shortest contigs we used only the largest contigs (>1Mb) for population genomic analyses. These large contigs accounted for 81% of the estimated genome size.

Protein-coding gene annotation was performed as previously described ^95^. Briefly, de novo and genome-guided transcriptome assembly was carried out using Trinity ^96^, followed by PSyTrans (https://github.com/sylvainforet/psytrans) to remove Symbiodinium transcripts. Transcripts were then assembled to the genome assembly using PASA ^97^, from which a set of likely ORFs were generated. Based on their protein coding ability and completeness, these ORFs were carefully assessed to produce a highly confident and non-redundant training gene set. This was used to train AUGUSTUS ^98^ and SNAP ^99^, and the resulting parameters were employed by the corresponding program in the MAKER2 pipeline ^100^, from which an ab initio gene model was predicted. Finally, putative transposable elements in the gene models were excluded based on transposonPSI (http://transposonpsi.sourceforge.net) and hhblits ^101^ search to transposon databases.

Functional annotation of gene models was performed as follows. For each gene in the *A. tenuis* genome, homologues with high quality functional annotations were identified using BLAST searches (E-value threshold 1×10^−5^) of predicted proteins (BLASTp) and predicted transcripts (BLASTx) against the SwissProt (June 2018) database. Genes were also annotated by searching for conserved domains using InterProScan^102^ (version 5.36-75). In addition to finding conserved domains, InterProScan results were also used to infer Gene Ontology based on the conserved domains present.

### Synteny Analysis

To infer macro-syntenic linkages, one-to-one mutual best hit orthologs were computed against *Nematostella vectensis* proteome (see section on orthology analysis for the list of all proteomes) using BLASTP. In total, 7,053 orthologous groups containing representatives from all 7 species were identified. Only scaffolds/chromosomes containing 10 or more such orthologous groups were considered. Macro-synteny plots were generated using a custom R script, where the synteny between *A. millepora* chromosomes to *A. tenuis* and *N. vectensis* scaffolds is represented by a dot for each orthologous gene family. The clustering of scaffolds for *A. tenuis* and *N. vectensis* was done by taking and sorting by the median position of the orthologous genes on *A. millepora* chromosomes.

### Orthology Analysis

Orthofinder^103^ was used to search for orthologous groups of genes shared between gene sets from the following cnidarian genomes, *Acropora digitifera* (Genbank GCF_000222465.1), *Acropora tenuis* (http://aten.reefgenomics.org/), *Acropora millepora ^45^*, *Fungia sp* (http://ffun.reefgenomics.org/), *Galaxea fascicularis* (http://gfas.reefgenomics.org/), *Nematostella vectensis* (Genbank GCF_000209225.1) and *Stylophora pistilata (*Genbank GCF_002571385.1).

### Sampling Design and Rationale for Population Genomics

Water quality is a significant factor influencing the health of inshore coral reefs^63,104^ and is subject to intensive monitoring on the GBR^65^. To capture adaptation in the *A. tenuis* holobiont related to water quality we chose reef sites at contrasting ends of two water-quality gradients both of which are well characterised in terms of exposure to riverine flood plumes^37,38,65,105^. The southern gradient is primarily influenced by secondary plume waters from the Burdekin river which affects reefs to a diminishing extent heading northwards. Predicted numbers of wet-season days exposed are Magnetic Island >67%, Pandora Reef 33-67% and Pelorus Island <33%^65^. A similar trend is also seen in the number of occurrences of low salinity events (<30) between 2003 and 2010 which were, Magnetic Island (8/8), Pandora Reef (6) and Pelorus Island (2). The northern gradient captures the wet tropics region where multiple rivers influence water quality. Here the plume location, Dunk Island is highly exposed (>67% of wet-season days; 8/8 low salinity events between 2003 and 2010) to plume waters from the Tully and to a lesser extent the Herbert rivers^37,106^ while Fitzroy Island receives flood waters from the Johnstone and Russell-Mulgrave river catchments much less often (lower end of 33-67% of wet-season days; 1/8 occurrence of low salinity between 2003 and 2010).

### Sample collection, sequencing and variant calling

Samples were collected in early 2015 from a total of 148 adult corals taken from five inshore locations on the central Great Barrier Reef (see Figure 2) including Magnetic Island and four northern locations, Fitzroy Island, Dunk Island, Pandora Reef and Pelorus Island. Samples were taken by cutting a branch and then placing it in ethanol for storage. Most samples were collected on a voyage of the RV Cape Ferguson in February (Fitzroy Island 20/2/15, Dunk Island 21/2/15, Pandora Reef 22/2/15 and Magnetic Island (Geoffrey Bay) 23/2/15). Half the samples from Pelorus Island were collected on 22/2/15 with the remaining 15 samples collected from nearby Orpheus Island (Pioneer Bay) on 24/4/15. Sequencing libraries were prepared separately for each sample and then used to generate four pools for sequencing across four flow cells of a HiSeq 2500 instrument (Ramaciotti Center for Genomics) to obtain a total of 1.2 billion 100bp paired end reads. With the exception of two samples that were chosen for deep sequencing (one from Fitzroy Island and one from Magnetic Island), libraries were pooled evenly so as to obtain approximately 3x coverage per sample. All population sequencing data is available under ENA study accession number PRJEB37470

Raw reads for all samples were pre-processed and mapped against the *A. tenuis* genome using the GATK best-practices workflow as follows. Reads passing quality checks were converted to unmapped bam files with sample, lane and flowcell tags added as appropriate. Adapters were marked using Picard (v2.2.1; http://broadinstitute.github.io/picard), mapping performed using bwa mem (v0.7.13; ^107^) and PCR duplicates marked using Picard (v2.2.1). Variants were then called using Freebayes (v1.0.2-16; ^108^) with haplotype calling turned off and ignoring reads with mapping quality less than 30 or sites with read quality less than 20. The two high coverage (20x) samples were downsampled to 3x using samtools to ensure parity with other samples prior to variant calling.

To ensure that only high-quality variant sites and high-quality genotype calls were used for population genomic analyses we performed the following basic filtering steps on raw variant calls from Freebayes. In cases where additional filtering was required it is described under the relevant analysis section. Sites with Phred scaled site quality scores less than 30 were removed. Only biallelic sites were retained. Sites where the mean read depth per individual was less than one third or greater than twice the genome wide average (2.7x) after mapping were removed. Genotypes with Phred scaled genotype quality score less than 20 were set to missing and sites with more than 50% missing genotypes were removed. Sites with highly unbalanced allele read counts (Freebayes allele balance Phred scaled probability > 20) were removed. The mdust program was used to mark low complexity regions and sites within these were removed. SNP sites within 10 base pairs of an indel were removed. Indel sites with length greater than 50 base pairs were removed.

### Population structure

To determine whether samples could be divided into genetically distinct clusters we used the program PCAngst (0.973)^109^ which has been designed specifically to infer population structure and admixture from low-coverage data. As input to PCAngst we used genotype likelihoods at all variant sites after quality filtering (see above), and converted these to Beagle format with ANGSD (0.920)^110^. Individual samples were plotted as a function of the top two principal components based on an eigendecomposition of the genotype relationship matrix calculated by PCAngst (Figure S2). This revealed two distinct clusters corresponding to the Magnetic Island and northern populations which is consistent with the optimal cluster number of 2, also inferred by PCAngst. Admixture proportions for each sample are shown in Figure 2 and show nearly complete assignment to either northern or Magnetic Island populations for most samples. Three samples (MI-1-1, PI-1-16, DI-2-4) showed mixed ancestry and were potential North/Magnetic Island hybrids while PCA analysis revealed two other samples (MI-2-9, MI-1-16) from Magnetic Island that did not appear to belong to either population cluster. We excluded all potential hybrids as well as these two outlying individuals from selective sweep and population demographic analyses.

### Demographic history with PSMC′

Changes in effective population size through time were inferred based on the genome wide distribution of heterozygous sites in two individual colonies sequenced to high coverage, (Fitzroy Island FI-1-3, 18x; Magnetic Island MI-1-4 20x) using PSMC′ implemented in the software MSMC (v2.1.2)^52^. In order to avoid biases due to differences in callability of heterozygous sites across the genome we only used these two high coverage samples and also used the snpable (http://lh3lh3.users.sourceforge.net/snpable.shtml) program to mask all regions of the genome where it was not possible to obtain unambiguous read mappings. Variant calling was performed for deep sequenced samples using samtools (1.7) and bcftools (1.9) as these generate outputs in the format required by the bamCaller python script included as part of the msmc-tools package (https://github.com/stschiff/msmc-tools). This script supplements the callability mask (see above) with an additional mask to avoid regions where read coverage is too low (less than half the genome-wide mean) or too high (double the mean) indicating a problematic region such as a collapsed repeat. As a final quality-control measure only scaffolds larger than 1Mb (~80% of the genome) were included in MSMC analyses as excessive fragmentation has been shown to bias results ^111^.

A distribution of MSMC estimates was obtained by generating 100 bootstrap runs, each of which involved recombining a random subsampling of 500kb-sized chunks to produce 20 large scaffolds of length 10Mb. Raw outputs from MSMC were converted to real values using a mutation rate of 1.86e^−8^ events per base per generation and a generation time of 5 years which is consistent with the fast growth rate and relatively high turnover of Acroporid corals.

The neutral mutation rate used in the calculation above was estimated by exploiting the equivalence between mutation and substitution rates for well-separated lineages (Kimura 1983). The well-studied congeneric species *Acropora digitifera* was chosen for this comparison because its divergence time from *A. tenuis* has been estimated at 15 million years ^40^, a value that is sufficient to allow near complete lineage separation but small enough that the number of multiply mutated sites should be small. First, 5583 one-to-one orthologs shared by *A. digitifera*, *A. tenuis* and *A. millepora* were identified using ProteinOrtho (v5.16b^112^). Amino acid sequences for *A. tenuis* and *A. digitifera* for these orthologs were then aligned with MAFFT (7.407^113^) and converted to codon alignments using pal2nal (v14 ^114^). Synonymous substitution rates, dS were estimated for each ortholog pair using codeml from the PAML suite (4.9i ^115^). Since dS is a ratio, its values follow a log-normal distribution and we therefore chose the central value as the mean of log(dS). Finally, the neutral mutation rate, μ was estimated using a divergence time (T) of 15.5Mya, a generation time (G) of 5 years and the formula μ = G*dS/(2T).

### Demographic history with ∂a∂i

Historical population sizes inferred under PSMC′ are made under the assumption of a single population without migration and have limited power to resolve changes in the recent past. In order to address these issues, we used the ∂a∂i framework ^116^ to fit a variety of more complex demographic models (Supplementary Table S2) to the joint site frequency spectrum between Magnetic Island and northern populations. A total of six demographic models were chosen to explore the timing and completeness of separation, population size changes and possible secondary contact between the northern and Magnetic Island populations. To explore the relationship between model fit and complexity we included models ranging from a very simple island model without migration (3 parameters) to a complex model allowing for population sizes before and after population split and asymmetric migration (10 parameters).

To prepare inputs for ∂a∂i we created a python script, vcf2dadi (https://github.com/iracooke/atenuis_wgs_pub/blob/master/bin/vcf2dadi.py) to calculate allele frequencies at each site based on observed read counts for each allele encoded in the Freebayes generated vcf file. The script rejects sites with fewer than 50% called genotypes and calculates allele frequencies by using the ratio of total observations for each allele to infer the number of alleles of each type present among called samples. This method takes advantage of the high overall read depth across all samples to accurately impute allele frequencies for each population without using potentially biased low-coverage genotype calls. Prior to running the vcf2dadi script further filtering was performed on the vcf file to remove sites with minor allele count less than 2, and to thin sites to a physical separation distance of at least 1000bp. This produced a final dataset consisting of 266k SNPs in ∂a∂i format. These were imported to ∂a∂i and converted to a folded 2D site frequency spectrum and projected down from the original sample size of 56 x 240 alleles to 45 x 180. This resulted in a final dataset that included 15.5k segregating sites.

All ∂a∂i models were optimised in a hierarchical manner to minimise the possibility of obtaining best-fit parameters corresponding to a local optimum. All models used generic starting parameters to begin ten independent runs, where each run involved four successive rounds with the best-fit parameters from the previous round undergoing perturbation prior to being used as input for the next round.

### Signatures of selection

A genome-wide search for selective sweeps was performed using SweepFinder 2 ^58,117^ which calculates a test statistic for each site in the genome that reflects the likelihood of observing SNP data under the assumption of a sweep compared with a null model where no sweep has occurred. In order to prepare inputs for SweepFinder 2 we used the same vcf2dadi script described under the section “demographic history with ∂a∂i”, with an option to generate outputs in SweepFinder format. The site frequencies generated by this script were then used to generate a combined frequency spectrum across the entire genome which SweepFinder 2 uses as an empirical neutral model. Each population was then scanned for sweeps by calculating the SweepFinder test statistic on a genome-wide grid every 1000bp. To avoid treating adjacent loci as independent sweeps we then ran another python script sf2gff.py (https://github.com/iracooke/atenuis_wgs_pub/blob/master/bin/sf2gff.py) which identifies contiguous sweep regions and converts them to gff format. In this script a sweep region is deemed to start when the sweepfinder statistic rises above a value of 10 and ends when it falls below this value. Within the region the maximum value of the test statistic is taken to represent the strength of the sweep. This resulted in 5315 putative sweeps which were then subjected to significance testing (see below).

Since the SweepFinder test statistic is a composite likelihood it requires an appropriate null model to determine a significance threshold. In order to address this problem and also to control for potential false positives arising from demographic factors we used the program, ms ^118^ to generate simulated allele-frequency data for all demographic scenarios modelled using ∂a∂i as well as the demographic history inferred by MSMC analyses. Neutral-model data simulated with ms was subjected to SweepFinder 2 analysis to calculate a null distribution of the composite likelihood ratio statistic (CLR) for each demographic model. At a given threshold value, T, of the CLR it is then possible to calculate the false discovery rate as the ratio of the number of loci with (CLR>T) in the neutral model versus the Magnetic Island or northern population data respectively. For the northern population a threshold CLR value of 100 or greater resulted in an FDR of less than 10% under any of the demographic models except for ‘sym_mig_size’ which was not the best-fitting model and predicted a deep bottleneck followed by population expansion. For Magnetic Island no such threshold could be found indicating that the influence of neutral factors was too large to allow genuine sweeps to be detected. For this reason we did not proceed further with SweepFinder analyses using the Magnetic Island data.

For the northern population all sweeps with a CLR value greater than a conservative significance threshold of 100 were identified and Bedtools (2.7.1^119^) was used to find genes that lay within 1000bp of these.

### Folded vs unfolded frequency spectra

We used the folded site frequency spectrum (SFS) for both dadi and SweepFinder 2 analyses. Although both methods can benefit from provision of an unfolded SFS this requires an accurate determination of the ancestral state for each allele. This is a particular challenge for *A. tenuis* because other *Acropora* species that could potentially be used as an outgroup (*Acropora digitifera*, *Acropora millepora*) are separated by a relatively large evolutionary distance (15 Million years ^40^). With such distantly related outgroup taxa there is a high likelihood that the outgroup sequence will not represent the ancestral state, but will instead capture independently fixed mutations in each lineage. We therefore decided to conduct all analyses with folded SFS in this study, which reduces power, but avoids the risk of false inferences due to incorrect ancestral assignment.

### Population genetic statistics, Fst, Tajima’s D

In order to avoid biases due to inaccurate genotype calls in low-coverage data we calculated Fst and Tajima’s D using ANGSD ^110^ (version 0.938) which calculates these parameters without calling genotypes ^120^. All calculations were performed based on the 14 million variant sites identified after quality filtering on Freebayes calls (see section on variant calling).

### Mitochondrial Genomes

Host consensus mitochondrial genome sequences were obtained for each sample as follows. Reads were mapped against the published *Acropora tenuis* mitochondrial genome sequence^121^ (genbank accession AF338425) using bwa mem (v0.7.17). Mapped mitochondrial reads for all samples constituted at least 25x coverage and were used to call a consensus sequence for each sample using bcftools (v1.9). Full length consensus mitochondrial genome sequences for all samples were used to build a TCS ^122^ type haplotype network and visualised using the software, PopArt ^123^ (v1.7)

Mitochondrial genome sequences for photosymbionts were obtained in a similar way. Reads previously identified as not mapping to the host genome were mapped against the *Cladocopium goreaui* mitochondrial sequences available from the reefgenomics website (http://symbs.reefgenomics.org/download/). Mapped reads were then used to call a consensus sequence using bcftools in the same manner as for host mitogenomes. Overall mitogenome coverage was much lower and more variable for symbiont genomes than for host genomes reflecting a larger mitogenome sequence (55kb) and reduced number of reads. We therefore removed samples with fewer than 2500 reads (~5x coverage) to avoid obtaining consensus calls from very low coverage. The resulting mitogenome sequences from 107 remaining samples were loaded into Geneious Prime (2019.2.3) and only sites with no ambiguous bases for any sample were retained. The resulting 6170bp segment was used to construct a TC haplotype network shown in Figure 2 using PopArt ^123^ (v1.7).

## Supporting information

Sweepfinder 2 Loci

## Acknowledgements

We would like to thank Dr. Anthony Bellantuono, Dr. Camila Granados-Cifuentes and Ms. Katherine Dougan for early access to the *Durusdinium trenchii* genome.

This project was supported by a Queensland Government DSITIA Accelerate Partnerships award to the University of Queensland on behalf of the Australian Institute of Marine Science (AIMS), the Australian National University, Bioplatforms Australia, the Great Barrier Reef Foundation, the Great Barrier Reef Marine Park Authority, and James Cook University (2014). AIMS staff Johnston Davidson, Paul Costello and Kathy Morrow are thanked for help in coral collections and Lesa Peplow for help with DNA extractions. Dr Gergely Torda collected the colony used for genome assembly and Dr Tom Bridge provided advice on taxonomy.

This research/project was undertaken with the assistance of resources and services from the National Computational Infrastructure (NCI), which is supported by the Australian Government. The data used in this project was funded by the Great Barrier Reef Foundation’s Resilient Coral Reefs Successfully Adapting to Climate Change research and development program in collaboration with the Australian Government, Bioplatforms Australia through the National Collaborative Research Infrastructure Strategy (NCRIS), Rio Tinto and a family foundation.

The authors also acknowledge the work done by the Reef Future Genomics (ReFuGe) 2020 Consortium organised by the Great Barrier Reef Foundation.

## Supplementary Information

Detailed methods including shell scripts used to run command-line analyses and R code used to generate plots is available from https://github.com/iracooke/atenuis_wgs_pub

The Acropora tenuis genome assembly and gene models are available for download from http://aten.reefgenomics.org/

https://docs.google.com/document/d/1EQqTLdvePd5iJTfdy88UVoY8aOG-ssX23HNv7AvGh0g/edit?usp=sharing

