## Supplementary Tables and Figures for "Signatures of selection in the coral holobiont reveal complex adaptations to inshore environments driven by Holocene climate change"

### Genome Statistics

**Table S1:** Table of contiguity and completeness statistics for published coral genomes

| Statistic | *Acropora tenuis (V2)* | *Acropora tenuis (V1)* | *Acropora digitifera* | *Acropora millepora* | *Acropora millepora* (V2) | *Fungia sp.* | *Galaxea fascicularis* | *Stylophora pistilata* |
|---|---|---|---|---|---|---|---|---|
| Num scaffolds | 614 | 1848 | 2421 | 3869 | 859 | 7424 | 11269 | 5688 |
| Total size | 486Mb | 408Mb | 447Mb | 387Mb | 475Mb | 606 Mb | 334 Mb | 400 Mb |
| Longest scaffold | 13.5Mb | 4.4Mb | 2.5Mb | 3.8Mb | 39.4Mb | 1.8 Mb | 0.9 Mb | 3.0 Mb |
| Scaffold N50 | 2.8Mb | 1.2Mb | 0.5Mb | 0.5Mb | 19.8Mb | 0.3 Mb | 0.09 Mb | 0.5 Mb |
| GC% | 39.07 | 38.93 | 39.4 | 38.84 | 39.06 | 38.41 | 39.43 | 38.53 |
| BUSCO completeness* | 90 | 89.9 | 74.4 | 90.5 | 90.4 | 86.5 | 87 | 88 |

### Repeat Content

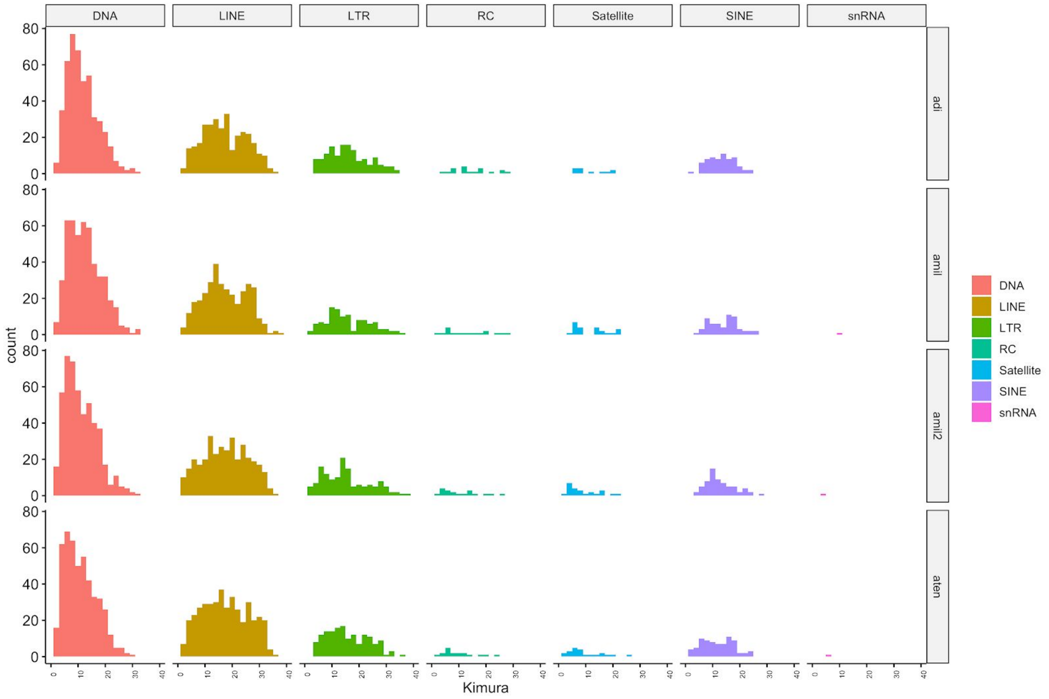


**Figure S1:** Relative proportion of major repeat classes in three available *Acropora* genomes (adi; *A. digitifera*, amil: *A. millepora*, amil2: *A. millepora* (long-reads), aten: *A. tenuis*). Counts of each type are plotted with respect to Kimura divergence.

### Coral Population Structure

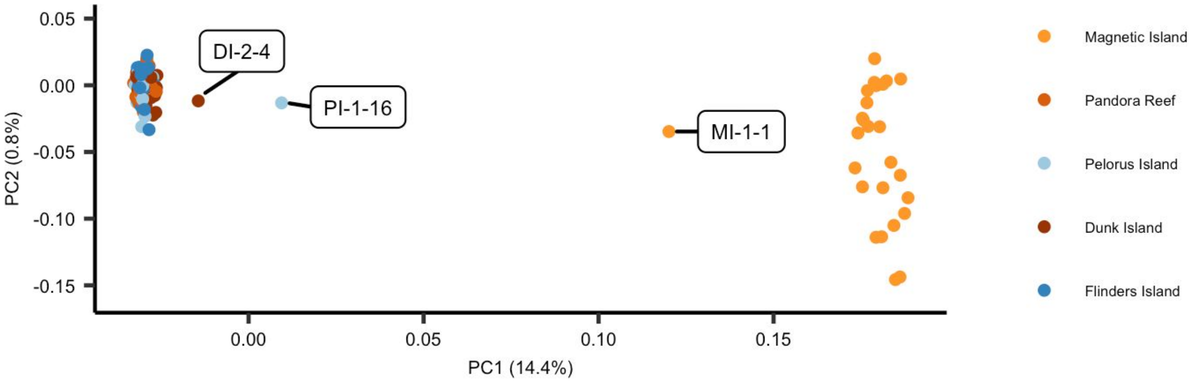


**Figure S2:** PCA plot showing coral host population structure separating northern sites from Magnetic Island. Labelled samples represent potential hybrids shown with stars in Figure 2B in the main text. Two outlying samples from Magnetic Island are not shown.

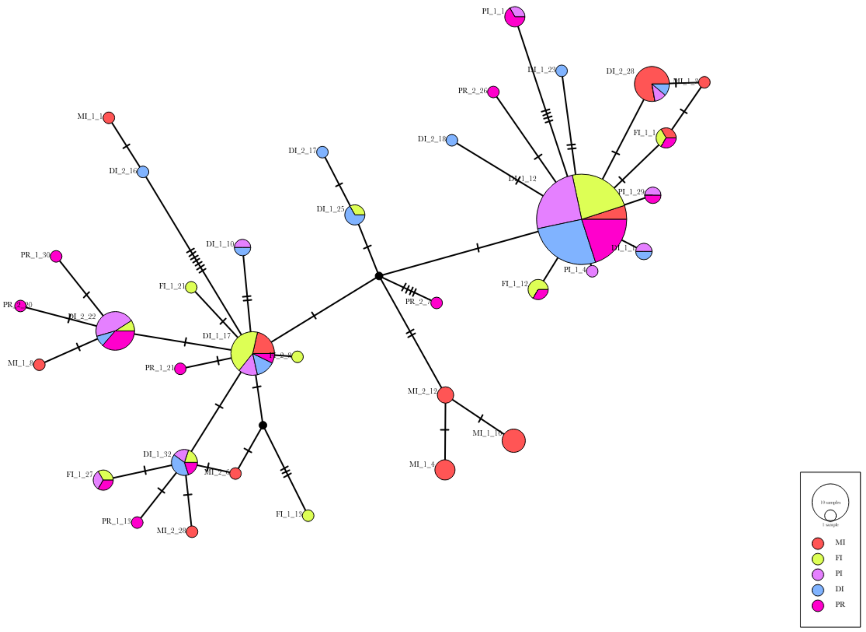


**Figure S3:** Mitochondrial haplotypes of *A. tenuis* represented as TCS network. Circle size represents the number of individuals sharing that haplotype with coloured segments showing the relative proportion from each of the five reefs. The number of lines on cross-bars between haplotypes represents the number of mutations separating them.

### Photosymbiont Profiles

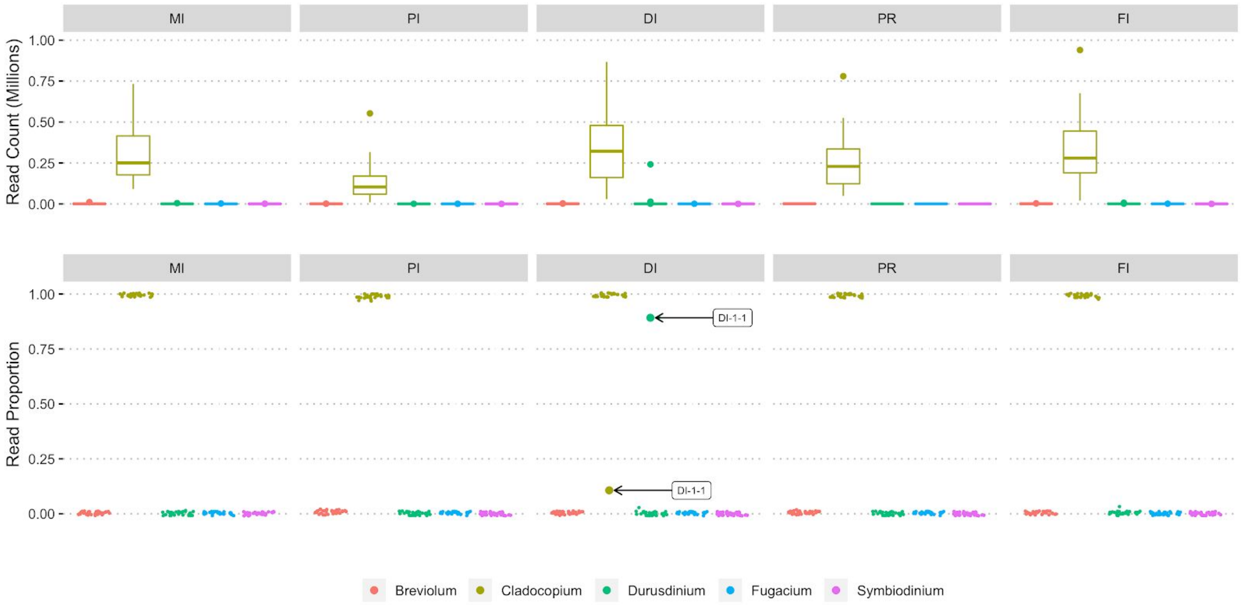


**Figure S4:** Symbiont profiles for all samples: (Top): Boxplots showing numbers of reads classified by genus for five genera in the family Symbiodinicaeae. (Bottom): Points showing the proportion of total symbiont classified reads in each sample. Most samples were exclusively dominated by reads from *Cladocopium* with the exception of a single sample (DI-1-1; highlighted with arrows) where the dominant symbiont was *Durusdinium*.

**Demographic Modelling**

**Table S2:** Demographic models and best-fit parameters from dadi analyses. Schematic images for each model show demographic history with blocks representing each of the two populations during a given time interval. Time is shown with the most recent at bottom and oldest at top. In all models population size is shown using parameters of the form $N^a_1$ where the superscript (a or b) represents the population (a=Magnetic Island, b = North) and subscript represents the time interval. Theta is the scaled mutation rate, $4N^{ref}\mu$. All times are given in years. Migration rate parameters are given in units of $2N^{ref}$ and are labelled as $m^{ab}$ to represent migration from b to a and vice versa. For the case of migration from b to a the value represents the proportion of individuals per generation in a that are due to influx from b.

| **Label** | no_mig | sym_mig | asym_mig |
|---|---|---|---|
| **Schematic** | 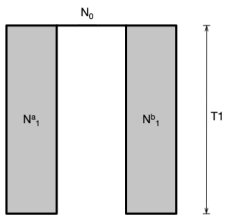  | 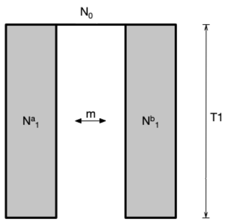  | 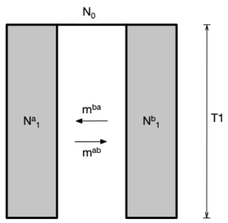  |
| **logLik** | -7854.74 | -6931.38 | -6650.19 |
| **theta** | 3471 | 2759 | 586 |
| **Divergence Time** | 4257 | 130,486 | 785,286 |
| **N0** | 96,771 | 76914 | 16,331 |
| **T1 (Years)** | 4257 | 130,486 | 785,286 |
| **$N^a_1$** | 3174 | 32,012 | 16,932 |
| **$N^b_1$** | 21,860 | 60,808 | 74,141 |
| **m ($m^{ab}$)** | | 1.1 | 3.2 |
| **$m^{ba}$** | | | 0.6 |

| **Label** | priorsize_asym_mig | asym_mig_size | isolation_asym_mig |
|---|---|---|---|
| **Schematic** | 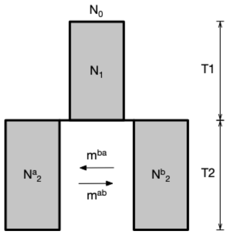 | 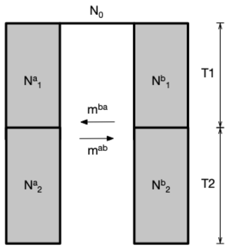 | 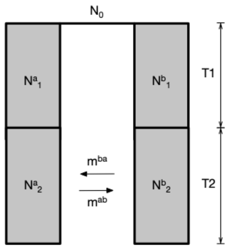 |
| **logLik** | -6318.77 | -6175.92 | -5855.16 |
| **theta** | 2286 | 29257 | 2240 |
| **Divergence Time** | 2071 | 254,881 | 268153 |
| **N0** | 63,735 | 815,620 | 62,448 |
| **T1 (Years)** | 93691 | 212,061 | 265530 |
| **N1 (Na1)** | 6,309,703 | 56,463,243 | 5,671,655 |
| **Nb1** | | 8,646 | 81,776 |
| **T2 (Years)** | 2071 | 42,820 | 2623 |
| **Na2** | 663 | 9624 | 681 |
| **Nb2** | 4621 | 64,271 | 12321 |
| **$m^{ab}$** | 92.4 | 90 | 95 |
| **$m^{ba}$** | 13.19 | 18.5 | 11 |

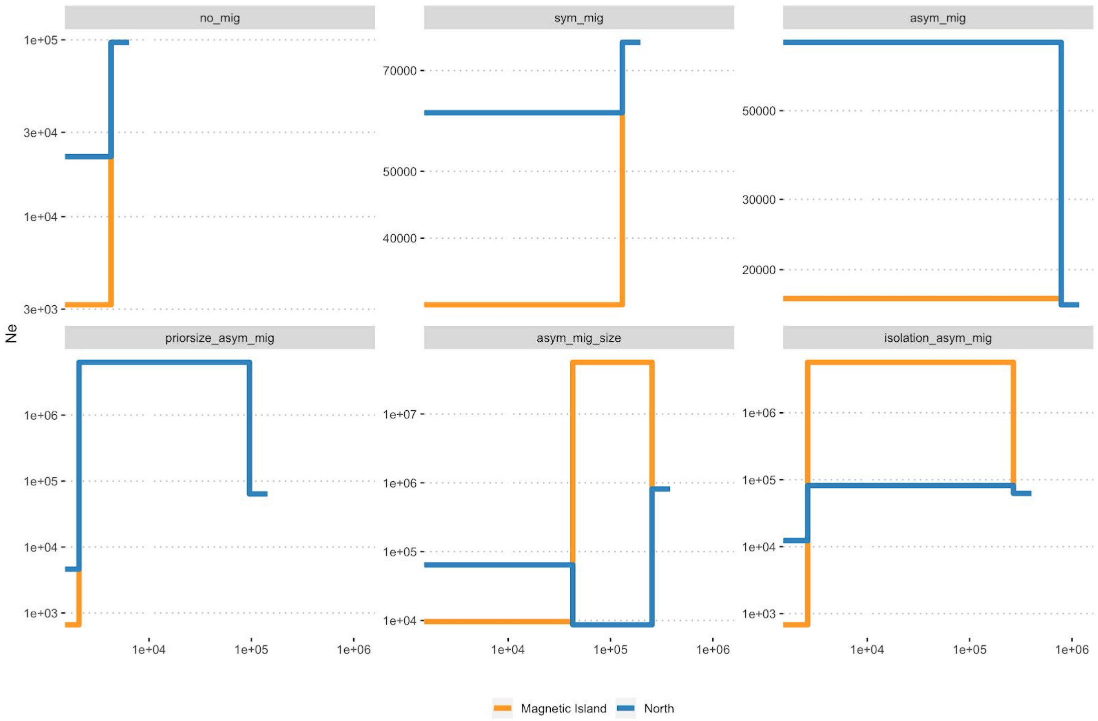


**Figure S5: Population size trajectories from all ∂a∂i models.** Each plot shows population size (vertical axis) trajectories for one of the models shown in Table S2 with Magnetic Island in orange and North in blue.  Time is shown with present day at the left and is in units of years before present. All plots use the same time axis but have different scales for population size.

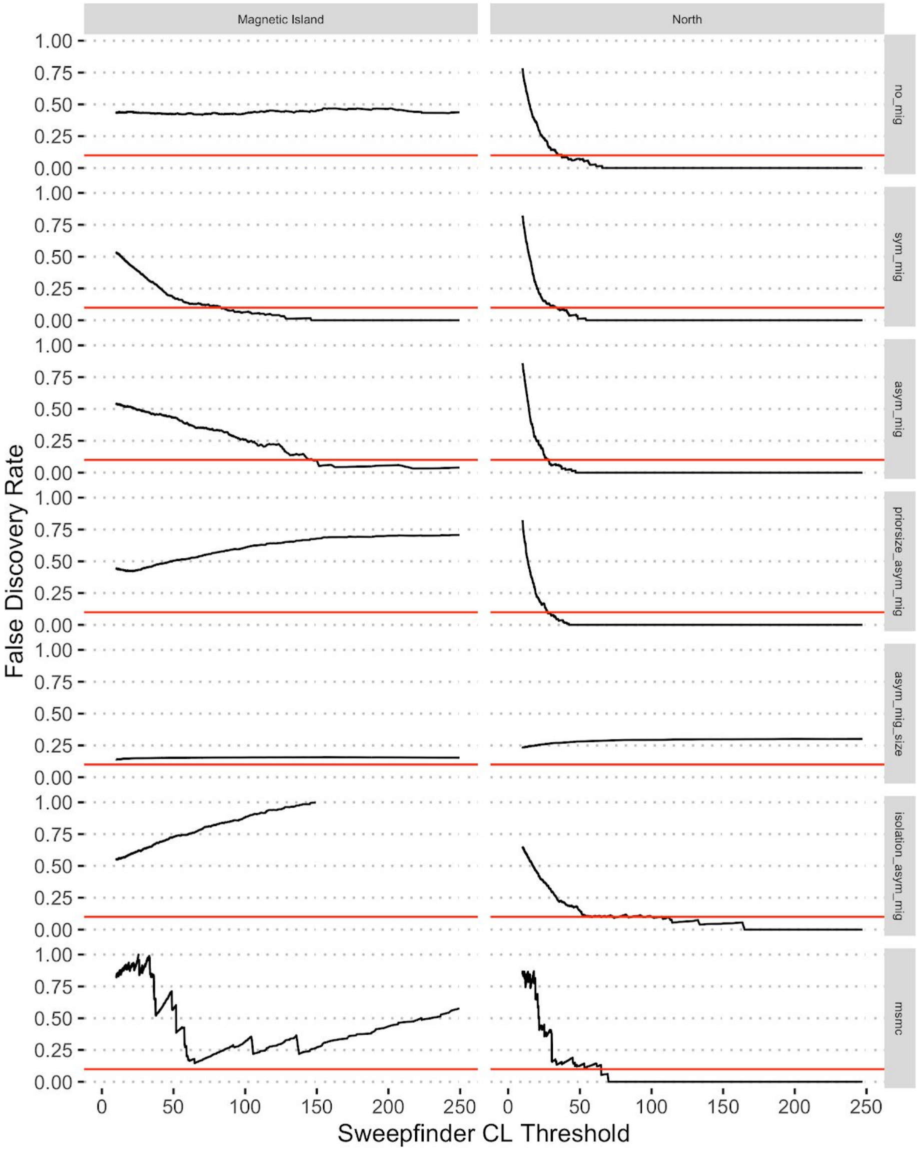


**Figure S6:** Empirical false discovery rate estimation for SweepFinder 2 statistics. Each sub-plot shows the calculated false discovery rate (FDR) as a function of SweepFinder score threshold for a different demographic model (see Table S2) and population. Red horizontal lines show an FDR of 10%.
